# The Origin of Human-infecting Avian Influenza A H6N1 Virus

**DOI:** 10.1101/000398

**Authors:** Liangsheng Zhang, Zhenguo Zhang

**Affiliations:** Department of Bioinformatics, School of Life Sciences and Technology, Tongji University, Shanghai, China; Advanced Institute of Translational Medicine, Tongji University, Shanghai, China; Institute of Molecular Evolutionary Genetics, Department of Biology, The Pennsylvania State University, University Park, Pennsylvania 16802, USA

## Abstract

In this study, we retraced the origin of the reported avian influenza A H6N1 virus infecting a 20-year-old woman in Taiwan. As we know, this is the first reported case of human infection by the H6N1 virus, because this subtype virus usually circulates in birds and poultry. Therefore it is crucial to know how this virus attained the ability to infect humans. Using phylogenetic analysis, we found that this virus was derived from reassortments of multiple lineages of H6N1 viruses and H5N2 viruses. The results deepen our understanding of how the new human-infecting virus originated and based on these we discussed possible explanations for the H6N1 infection of humans. Our results, together with recent studies of H7N9 viruses which result in severe disorders, suggest that reassortments among avian-type viruses are quite often, which may sometimes result in fatal infections in humans. Thus a close watch on the circulation of avian influenza viruses is pretty necessary.

## Results and Discussion

On June 21, 2013, Taiwan’s Centers for Disease Control (CDC) reported a pathogenic influenza A H6N1 virus^1^. Therefore it is crucial to understand how the capacity of infecting humans was attained. To approach this goal, we retrace the origin of this H6N1 strain through phylogenetic analysis.

The phylogenetic analysis of all H6N1 genes reveals four major patterns (Fig. 1A, and Fig. S1, materials and methods in Supplementary Appendix): (1) the pathogenic strain A/Taiwan/2/2013/H6N1 is always contiguous to the A/chicken/Taiwan/A2837/2013/H6N1 on the trees except for gene PB1; these two strains will be referred to as the 2013 H6N1 strains below; (2) the HA and NA genes of the 2013 H6N1 strains are most close to the H6N1 strains reported in years 2004/2005 (named 04/05 lineage) and to the H6N1 strains reported in 2009/2010 (09/10 lineage), respectively; (3) the PB1 genes of the pathogenic and chicken 2013 H6N1 strains are most close to the 04/05 lineage and the 09/10 lineage, respectively; (4) the other 5 internal genes of the 2013 H6N1 strains are contiguous to different H5N2 strains and then adjacent to the 09/10 lineage except for PA.

**Figure 1.**
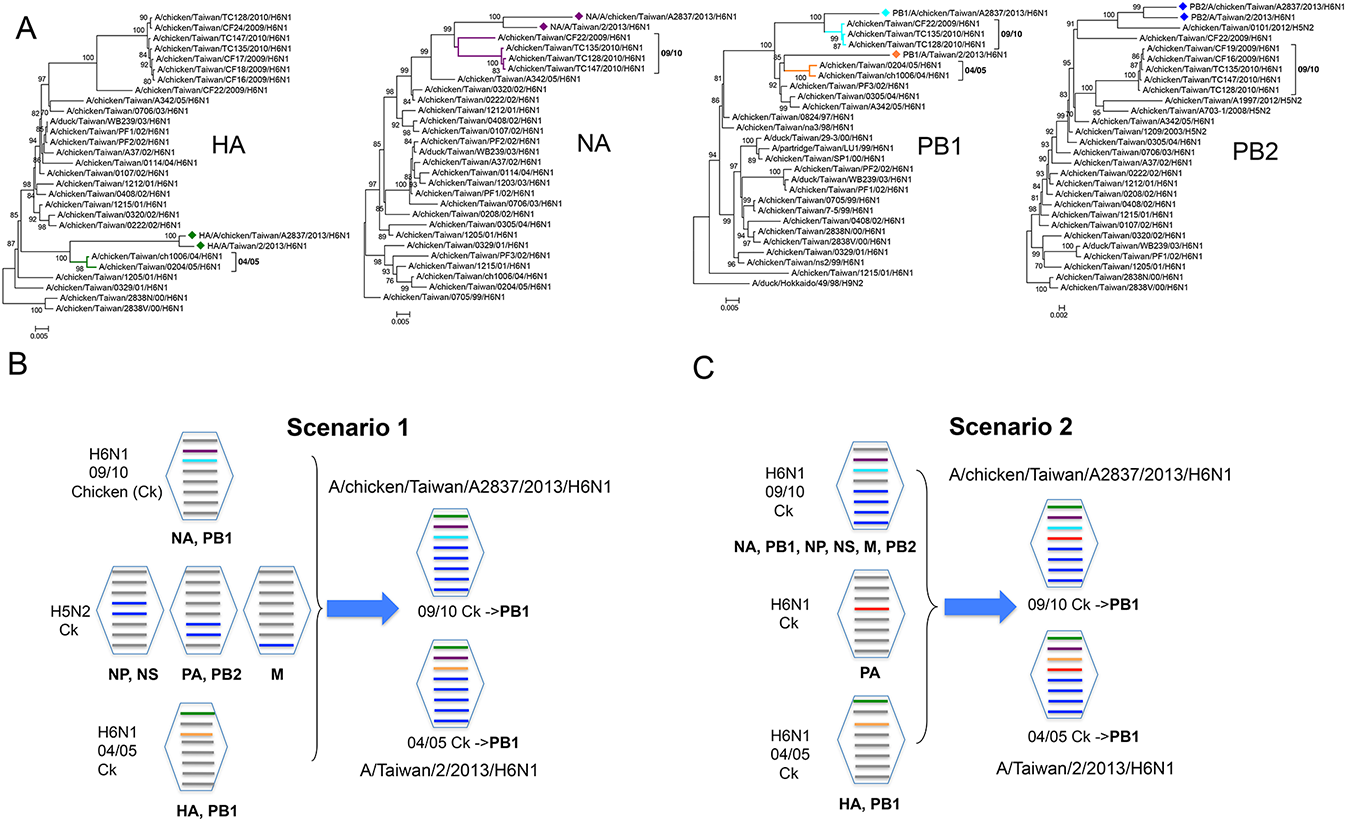
The phylogenetic trees of H6N1 genes and scenarios of reassortments generating the pathogenic influenza A (H6N1) virus. (A) Phylogenetic trees of HA, NA, PB1, and PB2 genes. The major lineages are labeled with colors. (B) and (C) The two proposed scenarios of reassortments to generate the two 2013 H6N1 viruses, A/Taiwan/2/2013/H6N1 and A/chicken/Taiwan/A2837/2013/H6N1. The 04/05 and 09/10 lineages are shown in (A).

According to these patterns, we propose two possible scenarios for the origin of the 2013 H6N1 strains. In the first scenario (Fig. 1B), the two strains may result from the reassortments of the 09/10, 04/05 and three H5N2 lineages. In the second (Fig. 1C), they may be from the reassortments of three H6N1 lineages while the neighboring H5N2 lineages in the trees of internal genes may be from multiple reassortments involving different H5N2 lineages and H6N1 lineage(s). To attest the two scenarios, more sequences of lineages near the common ancestral node of the H5N2 lineages and the 2013 H6N1 strains are needed. Yuan et al proposes that the 2013 H6N1 strains originated from a reassortment between H5N2 and H6N1 lineages^2^. The long branch between the 2013 H6N1 strains and the H5N2 lineages suggests that unknown evolutionary events may have occurred between the H5N2 to H6N1 viruses, and therefore, direct reassortment is unlikely to have occurred.

Based on our results, there are several possibilities for the explanation of the human H6N1 infection. (1) The reassortments to establish the two 2013 H6N1 lineages confer the viruses human-infecting capacity, but most infections are not detected because of low pathogenicity. (2) The mutations in the genes of the human H6N1 lineage after split from the chicken 2013 H6N1 lineage may be responsible for the new infection capacity. (3) Another possibility is that the patient might have some specific polymorphisms in the receptor genes, which make the binding of the H6N1 virus possible. These can be tested in future when data become available.

Our results, together with those of H7N9 studies ^2–5^, suggest that reassortments among different subtypes of avian influenza A viruses are quite common, which may sometimes generate pathogenic lineages. Therefore, a close eye should be kept on the surveillance of the avian virus circulations to prevent potential influenza epidemics.

## Acknowledgments

We acknowledge the authors, originating and submitting laboratories of the nucleotide sequences from Global Initiative on Sharing Avian Influenza Data (GISAID)’s EpiFlu Database (7/18/2013, 2 isolates). This work was supported by funds from Tongji University (985 programs).

### Conflict of Interest

All authors have no conflict of interest to declare.

